# Sound induces change in orientation preference of V1 neurons: Audio-visual cross-influence

**DOI:** 10.1101/269589

**Authors:** Nayan Chanauria, Vishal Bharmauria, Lyes Bachatene, Sarah Cattan, Jean Rouat, Stéphane Molotchnikoff

## Abstract

In the cortex, demarcated unimodal sensory regions often respond to unforeseen sensory stimuli and exhibit plasticity. The goal of the current investigation was to test evoked responses of primary visual cortex (V1) neurons when an adapting auditory stimulus is applied in isolation. Using extracellular recordings in anesthetized cats, we demonstrate that, unlike the prevailing observation of only slight modulations in the firing rates of the neurons, sound imposition in isolation entirely shifted the peaks of orientation tuning curves of neurons in both supra- and infragranular layers of V1. Our results suggest that neurons specific to either layer dynamically integrate features of sound and modify the organization of the orientation map of V1. Intriguingly, these experiments present novel findings that the mere presentation of a prolonged auditory stimulus may drastically recalibrate the tuning properties of the visual neurons and highlight the phenomenal neuroplasticity of V1 neurons.

**Highlights:**

- Prolonged application of sound shifts preferred orientation of neurons in area 17 of cats.
- Orientation tuning shifts in the individual neurons in supra- and infragranular layers of V1.
- Supra- and infragranular layers of V1 operate together but also as distinct compartments following sound adaptation.
- Visual cortex is specifically and dynamically impacted by non-visual stimuli.
- Sound adaptation certainly modifies the organization of orientation maps in V1.

## Introduction

For the longest time, inputs to a primary sensory area have been believed to be mainly modality specific. However, it was until recently (Ghazanfar & Schroeder, 2006; Kayser & Logothetis, 2007) that a direct and more specific interaction between the early sensory cortices of different modalities was discovered. Since then, more studies have suggested that the meeting and interaction of information from different senses begin within the low-level sensory areas (van Atteveldt *et al.*, 2014; Ten Oever *et al.*, 2015). Direct anatomical connections from V1 to A1 are still not precisely known, but across species there are different indirect pathways that have been discovered and suggest connectivity between different primary sensory regions. It has been shown in gerbils that secondary visual area V2 is directly connected with A1 (Budinger *et al.*, 2006). In cats, (Hall & Lomber, 2008) few direct projections from auditory cortex to V1 targeting only the peripheral visual field have been identified. An alternative yet slightly longer pathway between A1 and V1 through the heteromodal association cortical area (superior temporal polysensory area and superior temporal sulcus (STP ⁄ STS) was also revealed and presumed to send feedback to both A1 and V1 (Schroeder & Foxe, 2002). A critical imaging study (Liang *et al.*, 2013) revealed in humans that there might be salient locations within V1 that respond to specific cross-modal inputs related to the spatial pattern of activation of a primary sensory cortical area (e.g., V1).

Recently, more elaborate studies in rodents have exposed new findings. Physiological investigations of Ibrahim and co-authors (Ibrahim *et al.*, 2016) have demonstrated sharpening of orientation tuning in conjunction with an enhancement of the response to the preferred orientation of the cell in mice towards a low-contrast visual stimulus accompanied by an auditory stimulus. Lurilli and colleagues (Iurilli *et al.*, 2012) revealed that the presentation of a high-amplitude sound stimulus resulted in the hyperpolarization of the membrane potential of V1 neurons resulting in inhibition.

Despite extensive experimentation to explore wide-ranging interactions in the cortex, the understanding of cross-modal processing during the absence of direct stimulation of a corresponding sensory region remains implicit. To date, most studies have limited the presentation of stimuli for a very short duration. Additionally, rodents have been the first choice for electrophysiological experimentation in this direction and humans are limited to functional imaging techniques.

In the present investigation, a sound stimulus was applied to V1 neurons in cats in the complete absence of visual input for 12 minutes. Through extracellular recordings in area 17 of anesthetized cats, we found that in the presence of only a sound stimulus, the orientation tunings of simultaneously recorded layer 2/3 and layer 5/6 visual neurons were altered while exhibiting shifts in their tuning, that is a change of the preferred orientation. Also, the orientation selectivity shift magnitude was found to be larger in layer 5/6 neurons. We argue and suggest that the modification in the tuning properties was solely potentiated by the continuous repetition of the sound stimulus for 12 min and the spatial-temporal structure of the sound.

Interestingly, layer 2/3 neurons displayed an intriguing shift pattern towards horizontal orientations unlike cells of layer 5/6. Our data illustrate a novel extension to audio-visual cross-influence demonstrating that even an isolated application of a sound can robustly induce a reorganization of area V1.

## Experimental Procedures

### Ethical approval

Eight adult domestic cats (*Felis catus*) of either sex were used for experiments adhering to the guidelines approved by the NIH in the USA and the Canadian Council on Animal Care. Cats were supplied by the Division of Animal Resources of the University of Montreal. The animal surgery and electrophysiological recording procedures were performed according to the guidelines of the Canadian Council on Animal Care and were approved by the Institutional Animal Care and Use Committee of the University of Montreal (CDEA).

### Anesthesia

Cats were initially sedated with a mixture of Acepromazine (Atravet, 0.1 mg/kg, s.c., Wyeth-Ayerst, Guelph, ON, Canada) and atropine sulphate (Isopto Atropine, 0.04 mg/kg, s.c., Atrosa; Rafter, Calgary, AB, Canada) followed by a dose of anesthetic ketamine (Narketan, 40 mg/kg, i.m.; Vetoquinol, QC, Canada). Anesthesia was maintained during the surgery with isoflurane ventilation (2%, AErrane; Baxter, Toronto, ON, Canada). After the surgery, cats were fixed on the stereotaxic and were paralyzed by perfusion of gallamine triethiodide (Flaxedil, 40 mg/kg, i.v.; Sigma Chemical, St Louis, MO, USA). Artificial ventilation was maintained by a mixture of O_2_/N_2_O (30:70) and isoflurane (0.5%). The paralysis was continued by perfusion of gallamine triethiodide (10 mg/kg/h) in 5% dextrose lactated Ringer’s nutritive solution (i.v., Baxter, Mississauga, ON, Canada) throughout the experiment.

### Surgery

Local anesthetic xylocaine (2%; AstraZeneca, Mississauga, ON, Canada) was injected subcutaneously during the surgery before any opening of the skin. A heated pad was placed beneath the cat to maintain a body temperature of 37.5 °C. Antibiotics Trimethoprim and Sulfadiazine (Tribrissen; 30 mg/kg/day, subcutaneous; Schering Plough, Pointe-Claire, QC, Canada) and Benzylpenicillin procaine and Benzylpenicillin benzathine suspension (Duplocillin; 0.1 mL/kg, intra-muscular; Intervet, Withby, ON, Canada) were administered to the animals to prevent bacterial infection. First, a vein of the animal’s forelimb was cannulated. Then tracheotomy was performed to artificially ventilate the animal. A proper depth of anesthesia was ensured throughout the experiment by monitoring the EEG, the electrocardiogram, and the expired CO_2_:O_2_ saturation was kept in check using an Oximeter. The end-tidal CO_2_ partial pressure was kept constant between 25 and 30 mmHg. Third, craniotomy (1*1 cm square) was performed over the primary visual cortex (area 17/18, Horsley-Clarke coordinates P0-P6; L0-L6). The underlying dura was removed, and the multichannel electrode was positioned in area 17. The pupils were dilated with atropine sulfate (1%; Isopto-Atropine, Alcon, Mississauga, ON, Canada) and the nictitating membranes were retracted with phenylephrine hydrochloride (2.5%; Mydfrin, Alcon). Plano contact lenses with artificial pupils (5 mm diameter) were placed on the cat’s eyes to prevent the cornea from drying. Finally, at the end of the experiment, the cats were sacrificed with a lethal dose of pentobarbital sodium (100 mg/kg; Somnotol, MTC Pharmaceuticals, Cambridge, ON, Canada) by an intravenous injection.

### Stimuli and experimental design

Two types of stimuli were used - visual and audio. The receptive fields were located centrally within a 15° radius from the fovea. Monocular stimulation was performed. Receptive field edges (RF) were explored once clear detectable activity was obtained using a handheld ophthalmoscope (Barlow *et al.*, 1967). This was done by moving a light bar from the periphery toward the center until a response was evoked. Contrast and mean luminance were set at 80% and 40 cd/m^2^, respectively. Optimal spatial and temporal frequencies were set at 0.24 cycles/deg and in the range 1.0–2.0 Hz (at these values V1 neurons are driven maximally) for drifting sine-wave gratings (Bardy *et al.*, 2006). Then, gratings were presented randomly as visual stimuli covering the excitatory RF to compute the orientation tuning curves of neurons (Maffei *et al.*, 1973). Visual stimuli were generated with a VSG 2/5 graphics board (Cambridge Research Systems, Rochester, England) and displayed on a 21-inch Monitor (Sony GDM-F520 Trinitron, Tokyo, Japan) placed 57 cm from the cat’s eyes, with 1024×9×768 pixels, running at 100 Hz frame refresh (Figure 1A). The gratings moved unidirectionally in eight possible orientations presented randomly one by one. Each randomly presented oriented grating was given 25 times for 4s each with an inter-stimulus interval of 1-3s. Subsequently, the animal was exposed to broadband noise-like auditory stimuli comprising a range of frequencies. The auditory stimulus (3s 78 dB SPL) consisted of temporally orthogonal rippled combinations (TORC’s) with varying frequency components from 250 Hz to 8000 Hz (Fries *et al.*, 2007). The 3s stimulus was played continuously for 12 minutes, delivered by a pair of external loudspeakers, positioned perpendicularly relative to the animal’s axis at 57cm to the center of the fixation axis of the animal. The frequency response range of the speakers (range of audible frequencies the speaker can reproduce) was 120 Hz-18 KHz. For some recordings, the speakers were also displaced laterally at the same plane, at 30 cm on either side from the center of the fixation axis of the cat (Figure 1 A). This was done to test the change in responses if any when the position of the speakers was changed. The sound frequency and intensity were cyclical and optimized and set according to the experiment design using Bruel and Kjaer Spectris Group Sonometer. The stimulus was optimized on a standard C-scale of the sonometer for both ears. The spectrogram of the sound stimulus displaying varying frequencies is shown in Figure 1B. Immediately after the 12-min presentation of the acoustic stimulus, a series of drifting gratings was presented again in a random order (each oriented grating was presented 25 times where each of the 25 presentations was 4s and inter-stimulus time interval lasted 1-3s). It must be emphasized that sound was applied in isolation, that is, no visual stimulus was presented during the sound application. Finally, a recovery period of at least 90 minutes (Figure 1 C) was given for neurons to return to their optimal state as the recovery takes one log unit longer than adaptation duration (Dragoi *et al.*, 2000).

**Figure 1.**
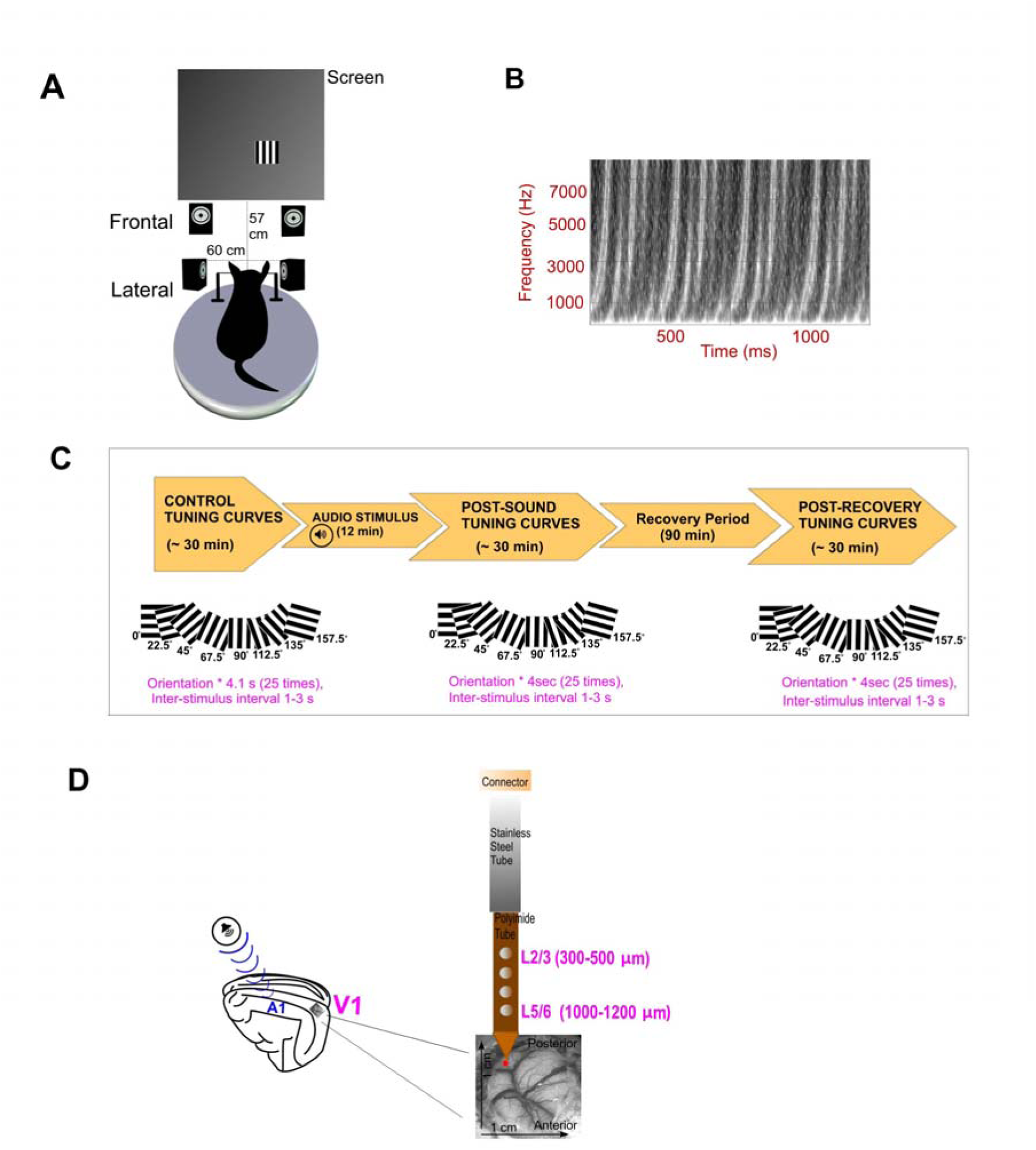
Experimental set up of stimulation, neuronal recordings and sound stimulus (A) Cartoon of the anesthetized cat fixed on the stereotaxic apparatus. Visual stimulus (shown as black and white gratings) was presented inside the receptive fields of test neurons. The sound stimulus was delivered by a pair of speakers occasionally placed frontal or lateral to the axis of the animal (B) The spectrogram of the sound stimulus is displayed. The spectrogram shows that sound stimulus was played in a constant frequency modulation (FM) pattern. (C) A pictorial representation of the steps followed during the protocol. Two types of stimuli were applied: Visual and auditory. Visual stimuli (sine-wave drifting oriented-gratings) were presented in a random order. Each orientation was presented 25 times and each trial lasted 4 s with a 1–3 ms inter-stimulus interval followed by the presentation of the sound stimulus for 12 min. Then, the same set of visual stimuli was presented again in a random order. A recovery period of at least 90 min was offered after which the gratings were presented again randomly. (D) Illustration of the electrode used to record neuronal activity. Neurons were recorded using multichannel electrode from depths 300-500 µm, and 1000-1200 µm from the primary visual cortex (V1)/area 17 before and after auditory stimulation.

### Electrophysiology

Multiunit activity in the primary visual cortex of anesthetized cats was recorded using a tungsten multichannel electrode (0.1–0.8 MΩ at 1 KHz; Alpha Omega Co. USA Inc). Neural activity was recorded from both hemispheres of the cat’s brain opening one side at a time. The electrode consisted of four microelectrodes enclosed in stainless-steel tubing in a linear array with an inter-electrode spacing of 500 µm. The recorded signal from the microelectrodes was acquired using Spike2, CED, Cambridge, UK. The signal was further amplified, band-pass filtered (300 Hz–3 KHz), digitized, displayed on an oscilloscope and recorded with a 0.05 ms temporal resolution. Recordings were performed at average cortical depths of 300–500 µm and 1000–1200 µm (Chanauria *et al.*, 2016) simultaneously from both sites as depicted in Figure 1 D. Spike sorting was done offline using same Spike2 package, CED, Cambridge, UK. Figure 2 shows an example of neuronal isolation (spike sorting) from the multiunit activity. As a precautionary measure, it was essential to affirm that we did not isolate the same unit twice, as the same unit may exhibit different waveforms depending upon several factors. Thus, the single units were distinguished based upon the spike waveforms, principal component analysis (PCA), and autocorrelograms (ACG) (Bharmauria *et al.*, 2015; Bharmauria *et al.*, 2016). The respective PCA, ACGs, and spike waveforms are also shown along in Figure 2.

**Figure 2.**
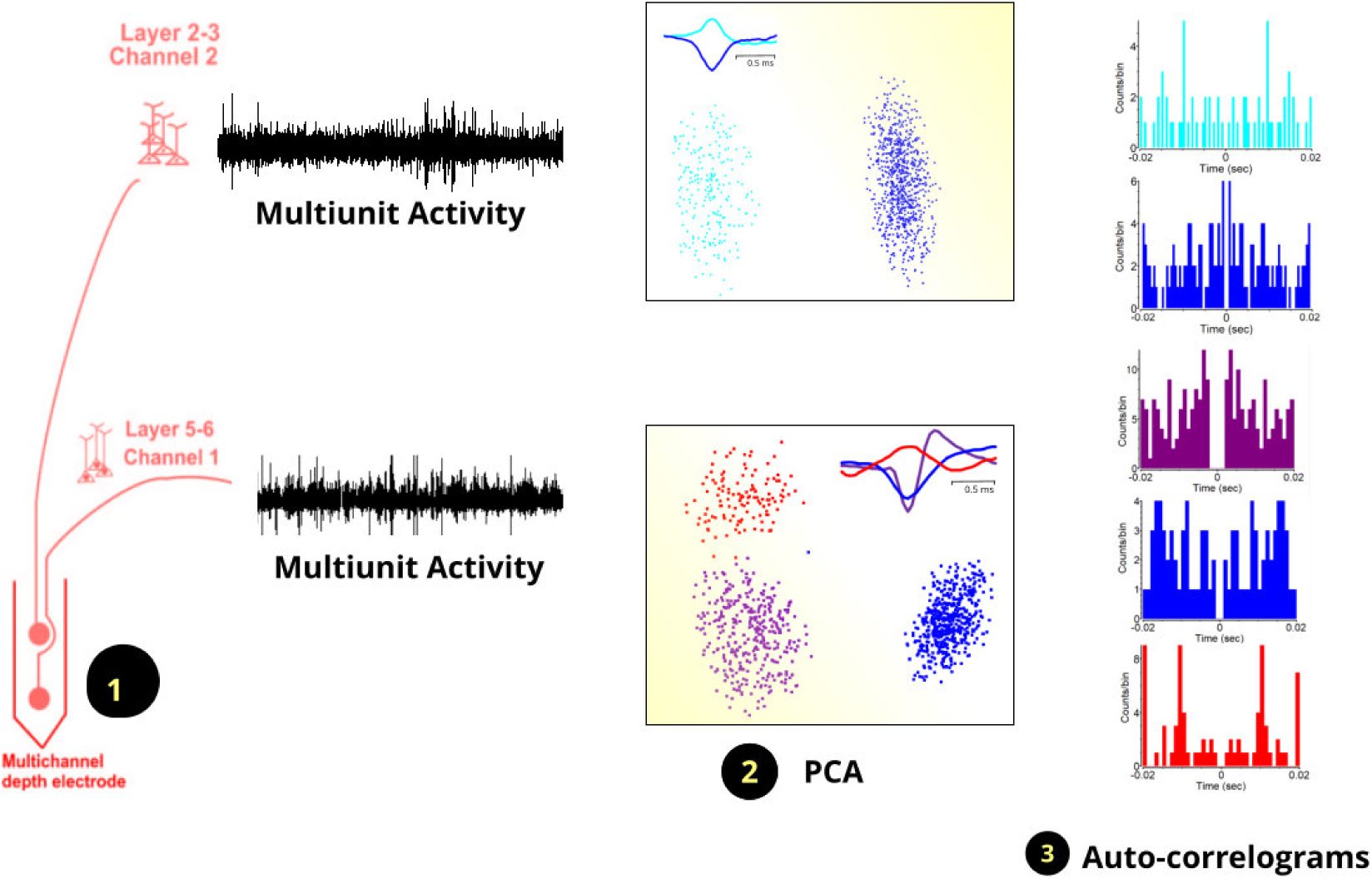
An example of the spike sorting process for isolation of single units is exhibited. Part 1 displays the neurons that have been recorded simultaneously from layer 2/3 and layer 5/6 of V1. Multiunit activity from both layers is also displayed alongside. Part 2 shows the principal component analysis of the dissociated waveforms. The superimposed average waveforms of dissociated spikes from multi-unit recordings are also shown. Part 3 displays the auto-correlograms for the separated single units. No events at zero represent the refractory period of neuron that confirms the individuality of each neuron.

## Data Analysis

### Tuning curves

After sorting neurons offline, the numerical value of orientation selectivity was calculated for each neuron at control and post-sound presentation conditions. Since applied gratings were separated by intervals of 22°, orientation peaks of tuning curves values were obtained by fitting a Gaussian non-linear curve on the raw orientation tuning values obtained at each orientation for each neuron. The Gaussian fits were calculated on trial-averaged data collected from 25 trials of the same stimulus at each orientation. Each point in Gaussian fits is the range of responses and thus represents the total response function. The Gaussian tuning fits were computed using the function below:

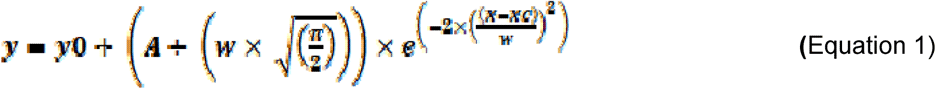

where *y0* is the offset, *XC* is the center, *w* is the width, and *A* represents the area under the Gaussian fit. The firing rates were normalized and gaussian tuning curves were generated in the scientific software Origin. The above procedure indicates the preferred orientation with better accuracy. The same method was carried in controls and following sound adaptation. The magnitude of shifts was computed as the distance (subtractions) between peak positions of the fitted Gaussian tuning curves before and after the presentation of sound (the difference between the initially preferred and newly acquired). The following formula was applied to calculate the shift magnitudes:

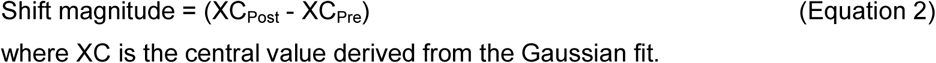

where XC is the central value derived from the Gaussian fit.

According to past studies, a difference of >5º is considered as a significant shift (Ghisovan *et al.*, 2009; Bachatene *et al.*, 2012; Bachatene *et al.*, 2013; Bachatene *et al.*, 2015; Chanauria *et al.*, 2016). A magnitude of <5º indicated a neuron that retained its initially preferred orientation even after the presentation of sound. These calculations are critical as the interval of 22° between the stimulus orientations is relatively large, which makes it difficult to deduce an exact value of orientation tuning from only raw curves.

### Orientation Selectivity Index (OSI)

Further, the Orientation Selectivity Index (OSI) of each neuron was computed by dividing the firing rate of the neuron at the orthogonal orientation by the firing rate of the same neuron at the preferred orientation, and subtracting the result from 1 (Ramoa *et al.*, 2001; Liao *et al.*, 2004). The closer the OSI is to 1, the stronger the orientation selectivity.

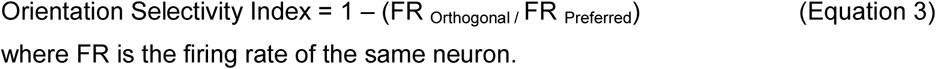

where FR is the firing rate of the same neuron.

### Bandwidths (BW)

Tuning bandwidths were calculated based on the full width at half magnitude (FWHM) of the Gaussian tuning curves for each neuron (Ringach *et al.*, 2002; Moore *et al.*, 2005). Bandwidths are measured to deduce the sharpness of orientation tuning curves of the neurons.

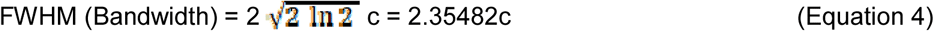

where ln is the logarithm and c is the gaussian root mean square width.

### Response Change Index (RCI)

To quantify the response disparity between stimulus conditions for each neuron, the traditional method of measuring response change index was used (Stevenson *et al.*, 2014; Meijer *et al.*, 2017) where the values of the RCI are normalized and can be used as a parameter to describe both enhancement and suppression. The values may range from −1 to 1 in which negative values indicate response suppression and positive values indicate response enhancement.

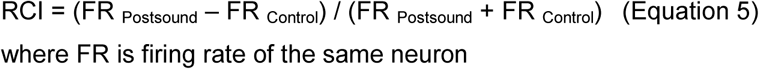

where FR is firing rate of the same neuron

## Statistical tests

Four datasets from layers 2/3 and 5/6 at control and postsound conditions were tested for statistics (i) orientation tuning values (ii) Bandwidth values (iii) Orientation Selectivity Index (OSI) values and (iv) Response Change Index (RCI) values. These data were first tested for normal distribution using the Shapiro Wilk normality test. Based on the results obtained, parametric and non-parametric tests were applied on data sets. Consequently, comparisons were drawn between values of different parameters for either layer. Detailed information of tests can be found in legends of the figures as well as the results.

The current investigation focused on how visual cells reacted towards oriented gratings when a sound stimulus was presented solely in the absence of any other visual stimulation. The extracellular activity of V1 neurons was simultaneously recorded from the layer 2/3 and 5/6 down the column. The Gaussian tuning curves of neurons in layer 2/3 and layer 5/6 were compared between the control and post-sound situations. In total, 239 cells were recorded during different experiments out of which 124 neurons belonged to the layer 2/3 and the remaining 115 to the layer 5/6. These pools were used for further analysis and statistics.

## Results

### Impact of repetitive auditory input on orientation tuning of visual neurons: A typical example

Figure 3 shows the typical result of a recorded neuron pair from layer 2/3 and layer 5/6. Two typical orientation tuning curves in raw forms are shown as (A) and (C) for layer 2/3 and layer 5/6 respectively. To infer the exact value of orientation preference, non-linear Gaussian fits were generated and are shown for the supra-granular (Figure 3B) and infra-granular cell (Figure 3D) for either condition. It must be emphasized that both cells were recorded simultaneously from the same electrode and the recording sites were separated by ~500 microns. The supra-granular cell exhibited an optimal orientation at 94.29° while the optimal orientation of the infra-granular layer was 96.70°, suggesting that the electrode was lowered in the same orientation column since optimal orientations are about equal. Following sound application, both cells displayed a novel optimal orientation. The peak of the optimal orientation of the supra-granular cell shifted to 110.92° indicating a displacement of 16° whereas the peak of the infra-granular cell moved in the opposite direction to 74.94° demonstrating shift amplitude of 21°. The opposite displacement of the peaks of optimal orientation tuning suggests that these shifts cannot be attributed to a global and spontaneous fluctuation of the firing rates. Each cell behaved independently. Furthermore, numerous studies have shown that, while the magnitude of the optimal responses may vary, the optimal orientation preference exhibits stability. Indeed, the optimal orientation remains the same for hours and even days and these controls were shown by Bachatene and colleagues (Henry *et al.*, 1973; Frenkel *et al.*, 2006; Lutcke *et al.*, 2013; Bachatene *et al.*, 2015). Therefore, the significant change in orientation selectivity is due to the experience of visual neurons with the sound. Unlike previous studies where a modulation of response was remarked after the presentation of auditory stimulus for a few milliseconds (Iurilli *et al.*, 2012; Ibrahim *et al.*, 2016), intriguingly, in this investigation, a complete shift in the curves of the orientation tuning of neurons was observed following the sound application.

**Figure 3.**
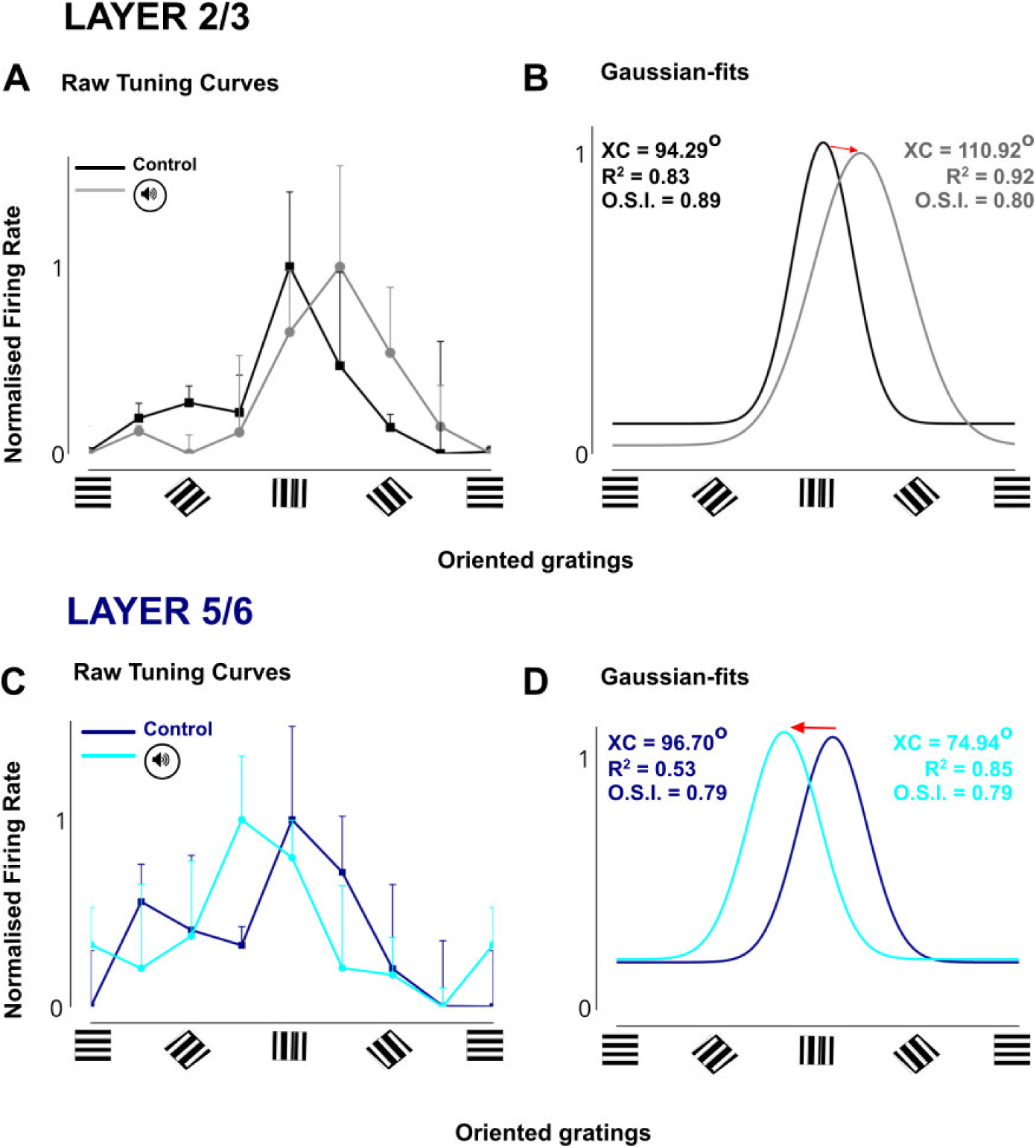
Typical examples of shift of orientation tuning peaks. The figure represents a pair of layer 2/3 and layer 5/6 neuron recorded simultaneously during the protocol. The bars signify standard error of the mean (mean ± SEM) over 25 trials at each presented oriented grating. The direction of shift is highlighted by an arrow (A) Raw orientation tuning values were compared for layer 2/3 neuron at control and post-sound conditions. A clear change in selectivity can be observed (B). To deduce the exact values of orientation tuning, raw values were fitted using the Gauss function. An overlap of the initially preferred and the new selectivity has been depicted along with the XC, R^2^ and OSI values. (C) Raw tuning curves of layer 5/6 neuron at control and post-sound conditions are displayed (D) Gaussian fitted tuning curves for layer 5/6 neuron over raw data is shown. The values of XC, R^2^ and OSI are also displayed alongside the curves.

### Neurons in supra- and infragranular layers regain their original optimal orientation tuning

Figure 4 shows examples of layer 2/3 and layer 5/6 neurons displaying a change in orientation selectivity during the presentation of oriented gratings at different steps of the recording protocol. Figure 4 A shows typical example of layer 2/3 neuron that was initially tuned to 82.06°. This indicated the optimal tuning of the neuron. Following the experience with 12 min of sound stimulus the same neuron changed its initially preferred orientation selectivity to a new preference of 32.80°. This neuron returned towards its original orientation selectivity, i.e., 78.01° after the rest period of 90 min. Likewise, the layer 5/6 neuron shown in Figure 4B recovered back from its new tuning peak of 115.73° to 91.09° after about 90 min of recovery time, which was quite close to its original optimal orientation tuning to 98.57°.

**Figure 4.**
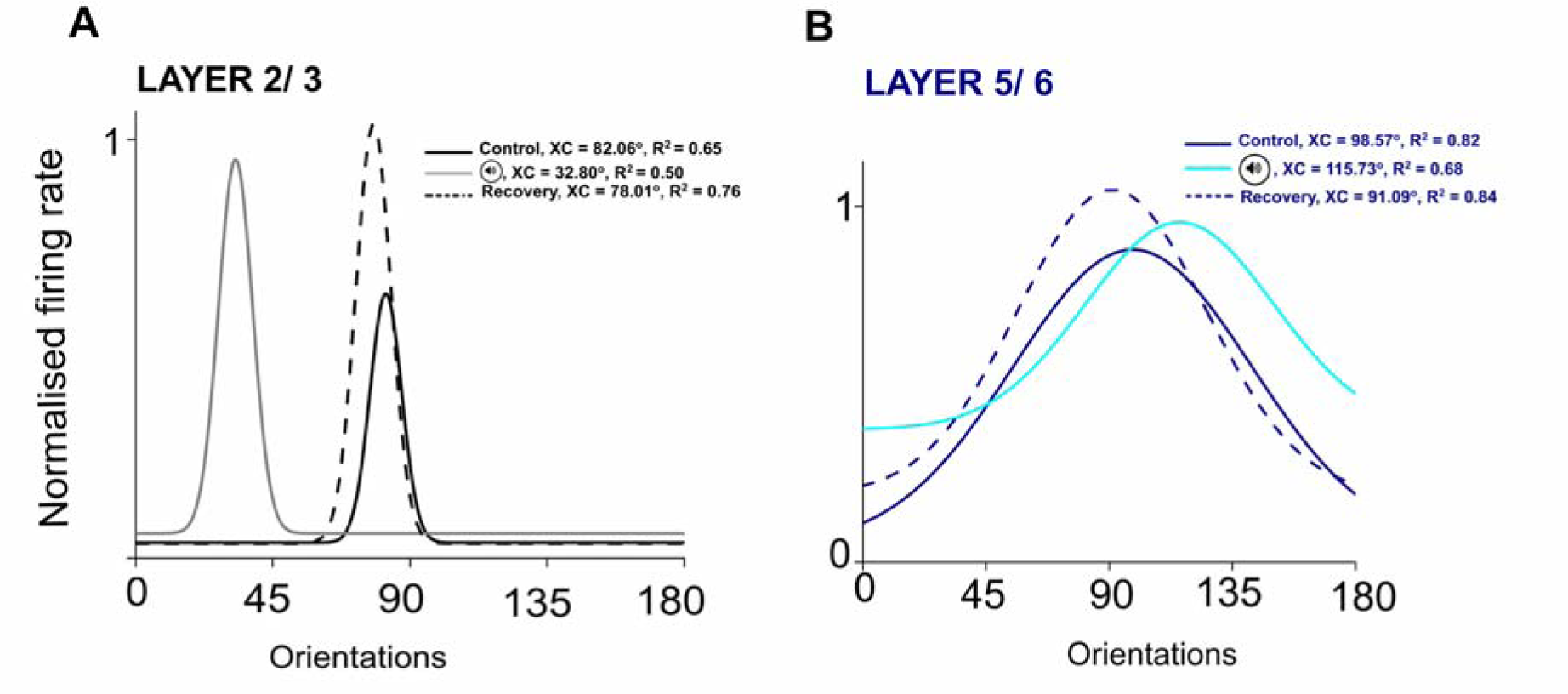
Typical examples of layer 2/3 and layer 5/6 neurons’ gaussian fits showing a change in orientation tuning curves at control, postsound and recovery stages. Both neurons clearly recovered back toward their original optimal after going through a resting period of 90 minutes. Figure 4A presents a layer 2/3 neuron that was optimally tuned to 82.06°. After 12 min of sound presentation the same neuron changed its orientation selectivity to 32.80°. Undergoing a rest period of 90 min, the same neuron returned towards its original orientation selectivity, i.e., 78.01°. Similarly, the layer 5/6 neuron shown in Figure 4B displays behavior of a neuron at different stages of sound adaptation protocol. The neuron was tuned to 98.57° initially, which was modified by the 12 min sound presentation to 115.73°. Finally, after the duration of 90 min, the same neuron gained its original orientation selectivity to 91.09°.

### Shift amplitude and sound source localization

The orientation tuning values of individual neurons at control and post-sound conditions are illustrated in Figure 5. On these datasets, a non-parametric statistical method was applied. The significant differences are indicated by solid black lines above each plot. The graphs A-C display values of orientation tuning peaks when speakers were placed laterally on either side of the animal with respect to the fixation axis of the animal. Figure 5A and 5B show the change in orientation tuning values pre- and postsound for layer 2/3 and layer 5/6 neurons, respectively. In Figure 5A (left side), on comparing the two conditions, the values were found to be significantly different. In Figure 5B (left side), the orientation preference values covered the full spread of presented gratings (range 0° to 157.5^0^). Remarkably, the postsound preferred orientations of layer 2/3 neurons appeared to have a significant bias towards horizontal orientations (range 0° to 50°), whereas this trend was not observed for cells in layer 5/6. The significant bias observed could be attributed to the variation of orientation selectivity by the properties of the auditory stimulus itself, or to the distinct mechanisms that drive the interaction between auditory and visual cortex. Further, different behaviors of neurons in different layers toward the same stimulus indicate different stages of cortical processing of the stimulus in the orientation column (Martinez *et al.*, 2005). The overall mean of the individual values (individual values and mean both shown) for the three experimental steps (control, postsound application, recovery ~90 min following sound application) are shown as inserts on the right side of Figure 5A and Figure 5B. There is a difference in number of neurons shown in these inserts. The recovery stage takes up about 90 min in the current study and sometimes even more. Only after this resting phase has passed the test orientations are recorded. Therefore, there arises a probability to lose the waveform of the neurons that were recorded during control and post-adaptation stages. It was of utmost importance to make sure of the waveforms at each of the three conditions. Therefore, only the neurons exhibiting similar waveforms at all three conditions were chosen.

**Figure 5.**
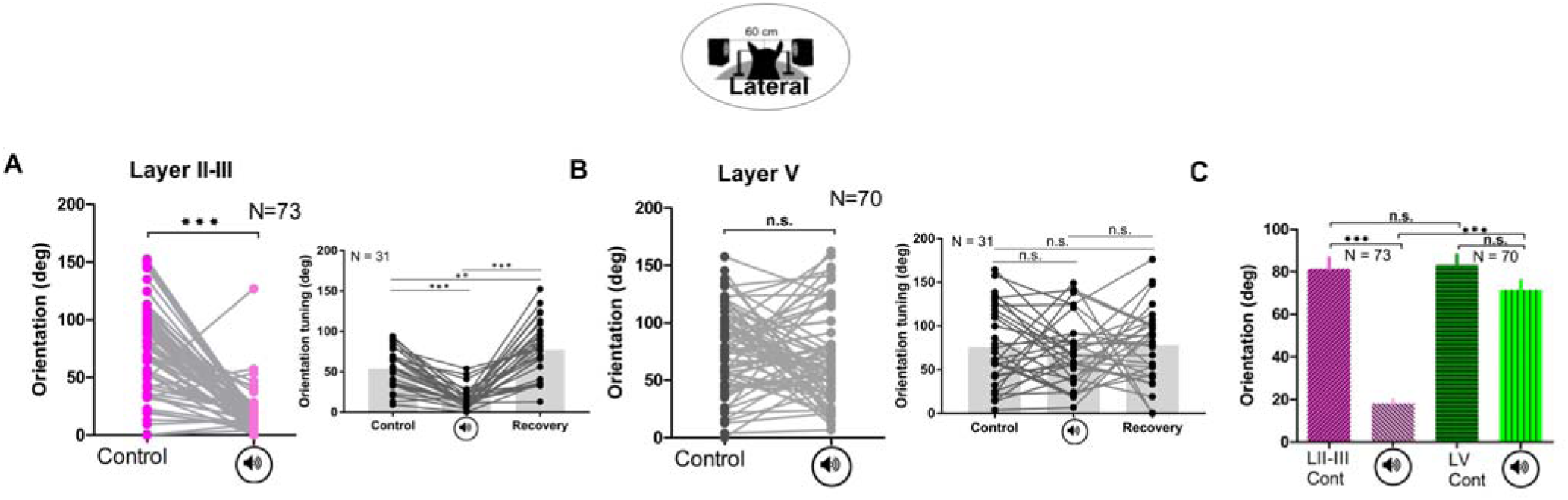
Acoustic stimulation results in modification of orientation preference in layer 2/3 and layer 5/6 neurons. (A) A plot of orientation tuning values for layer 2/3 neurons (N = 73), at control and post-sound presentation conditions is displayed (Wilcoxon matched-pairs signed rank test, P value <0.0001). On the right side, the histogram in grey and black shows the comparison of means of orientation selectivity values of (N = 31 out of 73) neurons at control, postsound and recovery conditions (Wilcoxon matched-pairs signed rank test: Cont-Postsound, P value <0.0001; Control-Recovery, P value <0.0047; Postsound-Recovery, P value <0.0001) (B) The graph shows the orientation tuning values for layer 5/6 neurons (N = 70) for control and post-sound conditions (Wilcoxon matched-pairs signed rank test, P value = 0.2280). The histogram in grey and black on the right side shows the comparison between means of orientation selectivity values (N = 31 out ot 70) at control, postsound and recovery conditions (Wilcoxon matched-pairs signed rank test: Cont-Postsound, P value = 0.6152; Control-Recovery, P value = 0.4487; Postsound-Recovery, P value = 0.8325) (C) This graph shows a comparison of means of orientation tuning values of layer 2/3 (N = 73) and layer 5/6 (N = 70) at control and postsound condition.

Further, mean orientation tuning values of layer 2/3 and 5/6 neurons at control and post-sound conditions were compared using Repeated measures ANOVA, P value = 0.0177 (refer to Figure 5C). The means (mean ± SEM) for the four mentioned data sets were 81.41 ± 4.952, 18.15 ± 1.882, 83.33 ± 4.566 and 71.38 ± 4.332, respectively. These four data sets were compared with one another using Tukey’s multiple post-comparison tests. Two out of six groups (layer 2/3 control versus layer 2/3 post-sound and layer 2/3 post-sound versus layer 5/6 control), were found to be significantly different, whereas the remaining four groups (layer 2/3 post-sound versus layer 5/6 post-sound, layer 2/3 control vs layer 5/6 control, layer 2/3 control vs layer 5/6 post-sound, and layer 5/6 control versus layer 5/6 post-sound) were found statistically non-significant.

To explore the effects of displacement of the sound source, the speakers were positioned in the front of the animal. Figure 6A and 6B showed the variation in displacements of orientation tuning for layer 2/3 (N = 08) and layer 5/6 neurons (N = 11), respectively, at control and postsound conditions. Layer 2/3 neurons showed a similar bias for horizontal orientations (tested significant statistically) in comparison to layer 5/6 cells. Figure 6C revealed the comparison of mean of orientation tunings of the above mentioned four groups. As an outcome of Repeated measures ANOVA, the comparison turned out to be non-significant. The means (mean ± SEM) for the mentioned four datasets were found to be 69.34 ± 13.21, 14.79 ± 4.90, 90.22 ± 11.91, and 91.24 ± 11.38, respectively (Figure 6 C). Using Tukey’s multiple post comparison test, two out of six groups; layer 2/3 post-sound vs layer 5/6 control and layer 2/3 post-sound vs layer 5/6 post-sound were statistically different whereas the remaining four groups layer 2/3 control vs layer 2/3 post-sound, layer 2/3 control vs layer 5/6 control, layer 2/3 control vs layer 5/6 post-sound and layer 5/6 control vs layer 5/6 post-sound were found statistically non-significant at the same confidence level. These results were similar to results obtained when the speakers were placed laterally on either side of the animal, thus revealing a lack of difference in responses of neurons. Together, these results displayed the extended plastic nature of layer 2/3 neurons over layer 5/6 neurons and indicated that the effects of sound stimulation are not impacted by displacement of the speakers.

**Figure 6.**
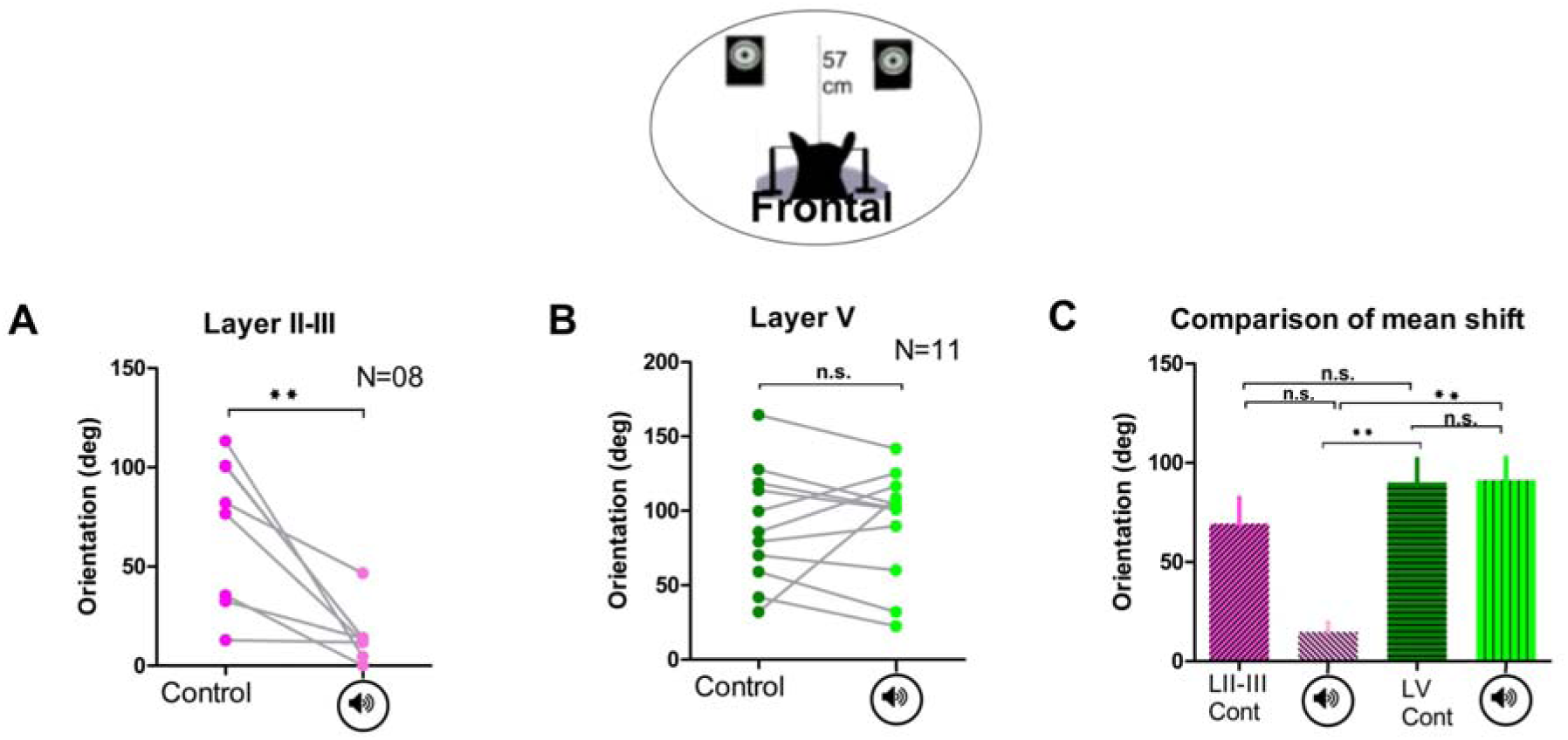
The graphs A-C show results when the speakers were positioned in front of the animal. A non-parametric statistical approach was followed since the data did not pass the Shapiro Wilk normality test. Connected points represent individual cells (A) Orientation tuning values for layer 2/3 neurons (N = 08) were plotted at both conditions and the difference was found to be significant (Wilcoxon matched-pairs signed rank test, P value = 0.0078) (B) Change in orientation tuning values of layer 5/6 neurons (N = 11) are displayed in the graph and the difference was found to be non-significant (Wilcoxon matched-pairs signed rank test, P-value = 0.8984) (C) Histogram of mean of amplitudes at control and post-sound conditions. The four groups (layer 2/3 control and post sound, and layer 5/6 control and post-sound) were compared using Repeated measures ANOVA. The means (mean ± SEM) for the four mentioned data sets were 69.34 ± 13.21, 14.79 ± 4.908, 90.22 ± 11.91 and 91.24 ± 11.38 respectively. Further, using Tukey’s multiple post comparison test, the differences between all groups were calculated. Two out of six groups, layer 2/3 postsound vs layer 5/6 control and layer 2/3 post-sound vs layer 5/6 postsound were statistically different, whereas the remaining four groups, layer 2/3 control vs layer 2/3 post-sound, layer 2/3 control vs layer 5/6 control, layer 2/3 control vs layer 5/6 post-sound, and layer 5/6 control vs layer 5/6 post-sound, were found statistically non-significant at the same confidence level.

### Sound impacts bandwidth of supra- and infra-granular neurons

The sharpness of orientation selectivity can be evaluated by measuring the bandwidth at half height of the orientation Gaussian tuning curve (Ringach *et al.*, 2002; Moore *et al.*, 2005). With this approach, we compared the tuning bandwidth of all neurons in either layer in control and post-sound conditions. The graphs shown in Figure 7 are a compilation of bandwidth analyses on layer 2/3 and layer 5/6 neurons. Figure 7A shows the difference in bandwidth for layer 2/3 neurons (N = 60). The large variance of the bandwidth of a neuron indicates broadening of selectivity for a range of orientations signifying a large contribution of flank orientations in the overall tuning of the neuron. Such an increase observed in our data strongly proposes that sound repetition induced the development of a different preferred orientation in the same neuron with a broader tuning curve. From this figure, overall, the tuning bandwidth at half magnitude (FWHM: full width at half magnitude) was found to be slightly enlarged at post-sound condition but not significantly different. This effect can be further understood by relating the results observed in Figure 4A where values of orientation preference for layer 2/3 cells held a bias towards the horizontal orientation’s postsound application. It gives the impression as if the neurons recorded from different sites in layer 2/3 were affected similarly on the application of sound and made all superficial layer neurons respond towards the sound in the same way. Similarly, in Figure 7B, layer 5/6 cells (N = 80) also showed a similar increment in the bandwidth responses. These were found to be statistically significant (Wilcoxon matched-pairs signed rank test, P-value = 0.0697). These results suggested that though both layers displayed similar behavior following the sound stimulus, nonetheless layer 5/6 cells exhibited broader shifts of tuning curves in comparison with layer 2/3 cells (refer to Figure 4B).

**Figure 7.**
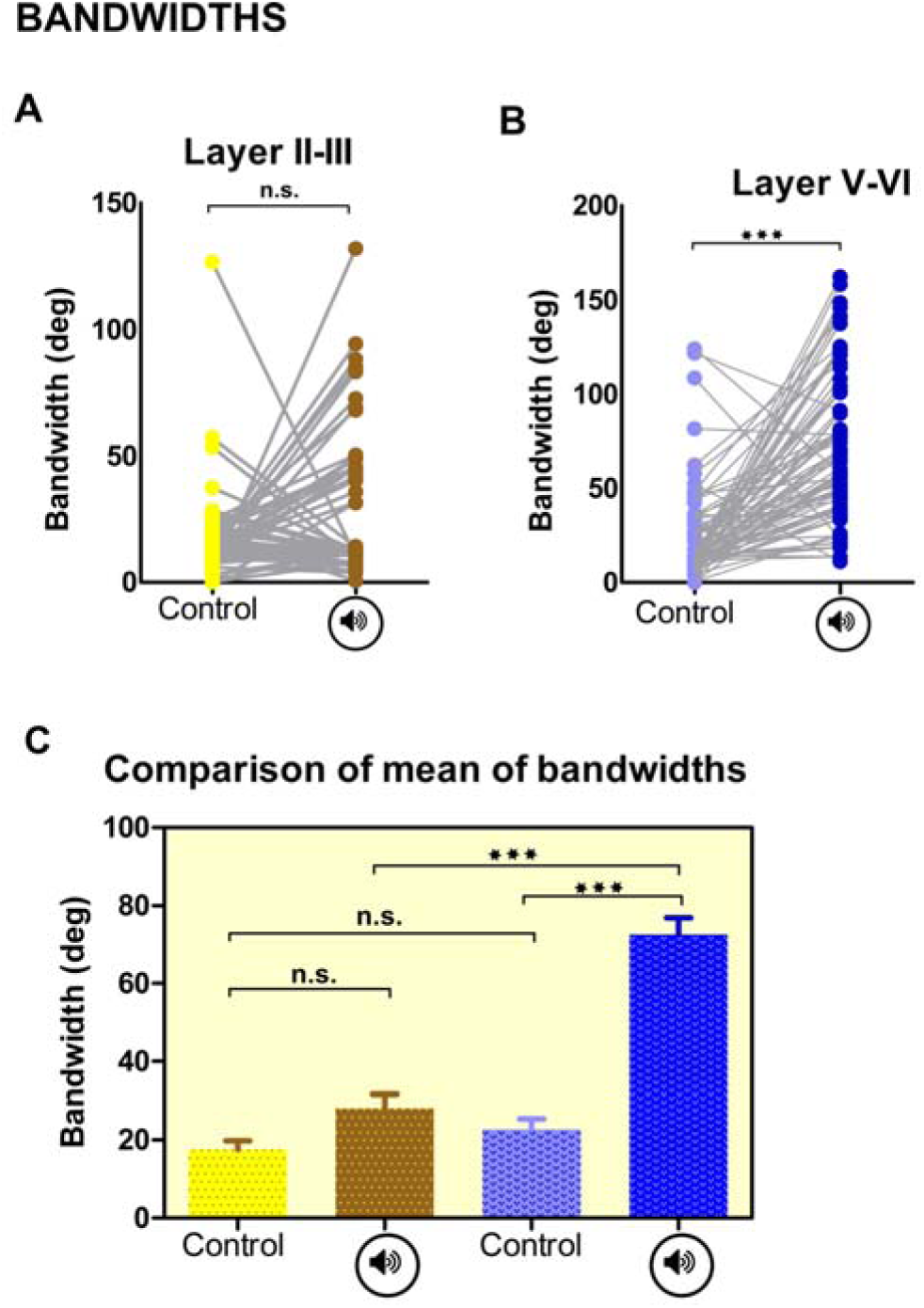
The sharpness of orientation selectivity. A non-parametric statistical approach was followed to measure significance between data sets. Connected points represent individual cells (A) The numerical values of bandwidths were plotted for layer 2/3 cells (N = 60) at control and post-sound conditions (Wilcoxon matched-pairs signed rank test, P-value = 0.0663) (B) A similar type of plot of bandwidth values for layer 5/6 neurons (N=80) is shown (Wilcoxon matched-pairs signed rank test, P-value = 0.0697). This group i.e. layer 5/6 group showed significant change in the bandwidth of neurons in comparison to layer 2/3 (C) Graph showing mean of bandwidths at control and post-sound condition. All four groups (layer 2/3 control, layer 2/3 post-sound, layer 5/6 control and layer 5/6 post-sound) were compared in this analysis using One-way ANOVA. The means (mean ± SEM) for the mentioned four data sets were 17.53 ± 2.293, 28.14 ± 3.595, 22.74 ± 2.736, and 72.52 ± 4.389 respectively. Further, with the help of Tukey’s multiple post comparison tests the differences between all four datasets were calculated. Three groups (layer 2/3 control vs layer 5/6 post-sound, layer 2/3 post-sound vs layer 5/6 post-sound and layer 5/6 control vs layer 5/6 post-sound) were found significantly different whereas remaining groups (layer 2/3 control vs layer 2/3 post-sound, layer 2/3 control vs layer 5/6 control and layer 2/3 post-sound vs layer 5/6 control) tested statistically non-significant at the same confidence level.

To test an overall trend, the mean values of neurons’ bandwidths were compared with each other in both layers (Figure 7C). All groups were compared with each other using Repeated measures ANOVA. The means (means ± SEM) for the mentioned four data sets were 17.53 ± 2.29, 28.14 ± 3.59, 22.74 ± 2.73 and 72.52 ± 4.38 respectively. Two out of six groups, (layer 2/3 post-sound vs layer 5/6 post-sound and layer 5/6 control vs layer 5/6 post-sound), were found to be significantly different (Tukey’s multiple post comparison test), whereas others (layer 2/3 control vs layer 2/3 post-sound and layer 2/3 control vs layer 5/6 control) were found statistically non-significant at the same confidence level. In summary, these results showed a global deviation of layer 5/6 cells with higher means of bandwidths.

### Orientation Selectivity Index (OSI)

In line with the previous reports (Dragoi *et al.*, 2001; Ringach *et al.*, 2002; Atallah *et al.*, 2012; Denman & Contreras, 2014) neurons having an OSI superior or equal to 0.4 can be classified as sharply tuned cells. In this investigation OSI values for all cells have been pooled to generate the figure. Figures 8A and 8B show scatter plots for regression analyses for layer 2/3 neurons (N = 124) and layer 5/6 neurons (N = 115) at control and postsound conditions, respectively. Layer 2/3 experienced more variation of response after sound application in comparison to layer 5/6. Overall, the OSI remained the same for either layer at both control and post-sound conditions yet layer 5/6 neurons experienced more dispersed evoked responses towards all presented orientations (Figure 8B). The comparison between all groups (using Tukey’s multiple post comparison tests) showed that layer 2/3 neurons experienced a decline in selectivity of the preferred orientation (Figure 8C). Two out of six groups (control layer 5/6 vs post-sound layer 2/3, control layer 2/3 vs post-sound layer 2/3) were found to be significantly different, whereas the remaining four groups’ differences were found to be non-significant. These results also correspond to our results from bandwidth data. The broadening of tuning curves after the imposition of sound lead to a decrease in mean OSI.

**Figure 8.**
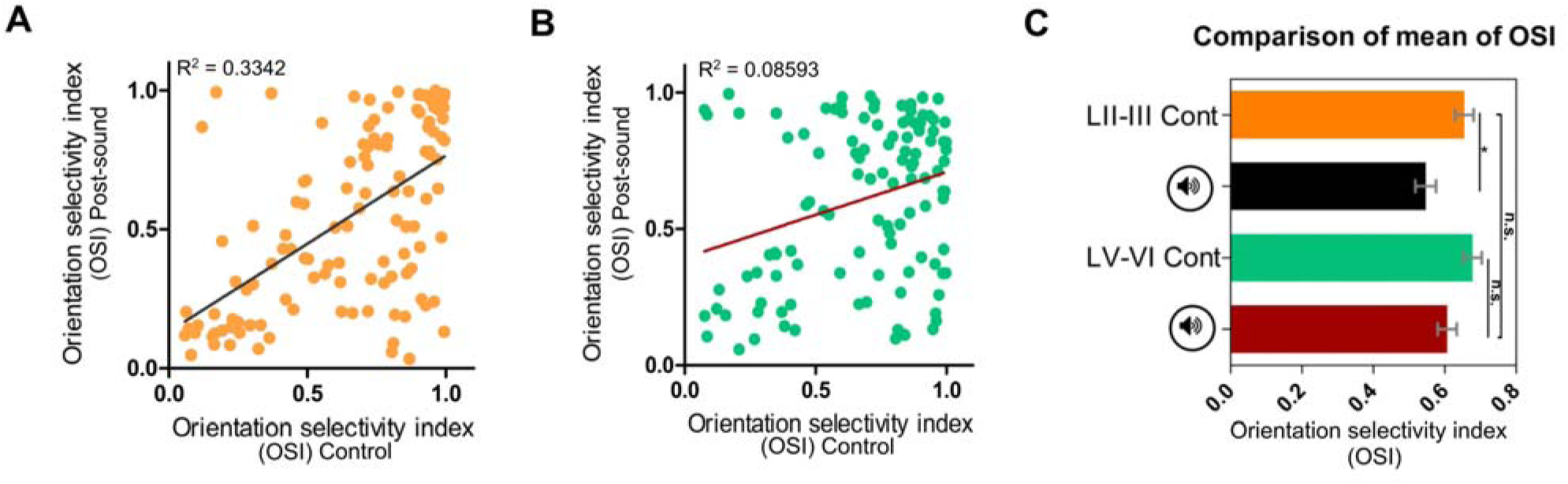
Orientation Selectivity Index (OSI) (A) OSI values calculated for layer 2/3 neurons (N = 124) at control and post-sound conditions are displayed as scatter plots and the regression line has been plotted (P-value<0.001; R^2^ = 0.3342) (B) Similar plot with regression analysis is also shown for layer 5/6 neurons (N = 115) at both circumstances (P-value = 0.0015; R^2^ = 0.08593) (C) The values of OSI from either layer for both conditions was compared using Repeated measures ANOVA. Only two groups tested statistically different for means (control layer 5/6 vs post-sound layer 2/3, and post-sound layer 2/3 and control layer 2/3) whereas remaining groups were not found different at any significance level.

### Response change index (RCI): Response modulation comparison between orientations

As described in the methods section the modulation of response magnitude was calculated for all cells and for all applied oriented gratings. The values for layer 2/3 (N = 124) and layer 5/6 (N = 115) cells have been pooled to generate the analyses. Results are compiled in Figure 9. The means RCI (mean ± SEM) of all layer 2/3 and layer 5/6 neurons (Figure 9A) were −0.03435 ± 0.02 and 0.03098 ± 0.02 and were found statistically non-significant (Mann Whitney Test, P-value = 0.0682). In layer 2/3, responses fluctuated roughly with the same magnitude for all tested orientations. The differences between all possible pairs among all orientations were measured by Repeated measures ANOVA and Tukey’s multiple comparison tests was applied to compare variances among means. The differences between all orientations in layer 2/3 were found to be non-significant (Figure 9 B). Interestingly, in layer 5/6 (Figure 9 C) the largest RCI was determined for orientations close to the vertical axis (recall cells in both layers were recorded simultaneously). A closer analysis indicates that RCI regularly declined to become negligible in both directions as evoked responses moved away from vertical to horizontal orientations. Only five (0° vs 90°, 22° vs 90°, 45° vs 90°, 90 vs 135° and 90° vs 157°) out of twenty-eight groups tested were significantly different. This analysis, however, unveils another important observation. When RCI was analyzed at individual orientations, we found that, in supragranular layers, the firing rate decreased for most orientations whereas in infragranular an opposite effect was observed. Nevertheless, the overall response variation is balanced (refer to Figure 9A). Altogether, layer 2/3 neurons experienced response suppression and layer 5/6 neurons experienced response enhancement.

**Figure 9.**
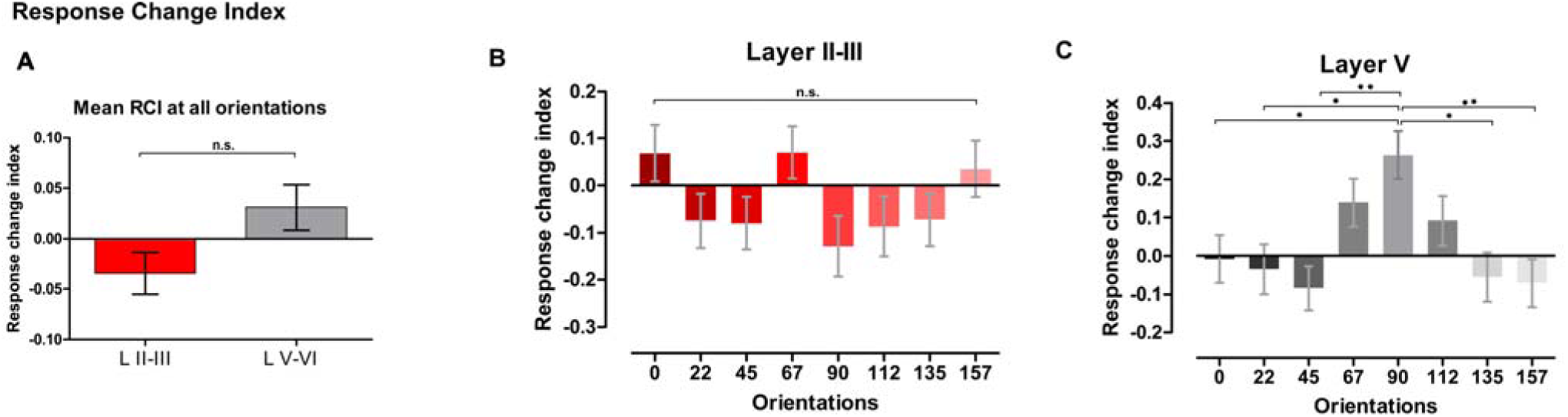
Response Change Index (RCI). A non-parametric method was opted for statistical validations (A) Histogram showing mean of RCI of all test neurons in either layer at all presented orientations (0° − 157.5°, separated by 22. 5°). The mean RCI (mean ± SEM) of all layer 2/3 and layer 5/6 neurons were − 0.03435 ± 0.02054 and 0.03098 ± 0.02241 and statistically non-significant (Mann Whitney Test, P-value = 0.0682) (B) A graph showing histograms of RCI at different orientations for layer 2/3 cells. The differences between all possible pairs among all orientations were measured by Repeated measures ANOVA and Tukey’s multiple comparison test. (C) A similar analysis was done on layer 5/6 neurons, and the histogram was generated. Five groups out of all possible paired comparisons (0° vs 90°, 22° vs 90°, 45° vs 90°, 90° vs 135°and 90° vs 157.5°) tested significantly different, whereas, for rest of the groups, the mean of RCI was not significantly different.

Finally, the striking inverse response modification of layer 2/3 and layer 5/6 neurons further suggests that neurons in supra- and infra-granular layers behave differently towards the identical stimuli conditions and it appears cells of both layers behave independently to process information. This kind of behavior where layers conduct themselves differently towards the same stimulus is indicative of different stages of cortical processing in a particular column (Martinez *et al.*, 2005).

## Discussion

The present study aimed to investigate the responses of visual neurons of V1 when stimulated by a repetitive auditory stimulus whose frequency and power varied cyclically, presumably driving many putative synaptic inputs as auditory cells respond to many different sounds. The investigation revealed three main findings following adaptation protocol to sound. First, and the most important result, we observed shifts in the orientation tuning curves in cat’s supra -and infra-granular layer neurons implying new orientation selectivity. Second, superficial and deeper layer neurons, recorded from the same columns at the same time, seemed to behave differently.

Moreover, occasionally cells in both layers shifted in opposite directions indicating independent behavior. Third, neurons in infra-granular layer 5/6 exhibited larger shifts of their orientation selectivity than the layer 2/3 neurons. Collectively, these findings suggest that neurons in the primary visual cortex (V1) might have a supplementary role in addition to processing a visual stimulus. Surely, these observations unveil that neurons in the primary visual cortex may alter their orientation preference since the peak of the tuning curves points to a novel preferred orientation.

### Methodological considerations

We recognize that anesthesia affects the frequency of neuronal firing, especially when comparing anesthetized with behaving animals. Under anesthesia, neurons fire more sparsely and with lower firing rates (Barth & Poulet, 2012). Land (Land *et al.*, 2012) revealed the effect of isoflurane on the cortical processing in a cross-modal study of the visual and auditory cortices of mice. However, the effects observed in Land’s study were an outcome of a deliberate decrease and increase in the levels of isoflurane during the recording sessions. In the present case, anesthesia was kept constant. Also, several findings argue against a significant influence of anesthesia. In general, anesthesia decreases the neuronal responsiveness globally. Shifts in the orientation tuning curves are ascribed to a decrease of responses to preferred orientation and simultaneously an increase to non-preferred orientation, that is, a push-pull mechanism. Similar shifts in orientation selectivity were reported when the adapting stimuli were visual in many species (Dragoi *et al.*, 2000; Ghisovan *et al.*, 2008; 2009; Bachatene *et al.*, 2012; Jeyabalaratnam *et al.*, 2013; Chanauria *et al.*, 2016).

Furthermore, the shifts in supra- and infra-granular layers were in the opposite direction. Additionally, supra-granular cells exhibited a tendency to shift their axis of orientation toward horizontal while infra-granular cells shifted the preferred orientation over the entire spectrum (note that cells were recorded simultaneously with the same electrode, and orientations were applied in random order). In cats, it has been shown that orientation selectivity does not occur at the LGN (lateral geniculate nucleus) level but is initiated at the cortical level (Dragoi *et al.*, 2001; Bouchard *et al.*, 2008). Infra-granular cells exhibited response change indexes increase predominantly for orientations close to vertical. Thus, such differential effects are difficult to reconcile with the global effects of anesthesia.

It could be of concern as to why a stimulus of 12 min was chosen. The reason behind the choice was first, to stimulate the auditory cortex with varying frequencies for a prolonged duration that could further impact the visual cortex. The current literature showing effects of the auditory stimulus on visual neurons used short sounds that induced modulation of the firing rate. Thus, it became of interest to observe responses of visual neurons in response to a longer duration sound. Since it was already established that 12 min duration is enough to entirely change the tuning properties of neurons we used a sound stimulus for 12 min. The sound stimulus lasted 3s and was presented uninterruptedly for 12 min. Therefore, the duration of the sound stimulus was selected to compare the effects with visual adaptation (Chanauria *et al.*, 2016) and auditory adaptation on supra- and infra-granular visual neurons. It was of utmost interest to see if neurons behaved similarly or not by triggering the visual cortex by another modality. Of course, it remains crucial to examine the effect of sounds with different properties and different durations on the visual cortex; that will help better understand the interactions between auditory and visual cortex.

### Layers behave as separate compartments

Our results also show that the application of sound differentially affected the layers 2/3 and 5/6, thereby also reconfiguring the layout of orientation maps in the V1. Ibrahim and co-authors (Ibrahim *et al.*, 2016) demonstrated that layer 2/3 neurons’ responses are suppressed by the A1 signals. Layer 2/3 neurons keep receiving the information for 12 min and presumably the intensity of the inhibitory drive increases with the duration of adaptation. Interestingly, it has been suggested that a supplementary excitatory feedback drive initiated in A1 may interact with layer 5/6 neurons (Fritz *et al.*, 2003; Muckli & Petro, 2013; Vetter *et al.*, 2014; Muckli *et al.*, 2015; Ibrahim *et al.*, 2016). This dual projection of neurons in supra- and infra-granular layers may elucidate shifts of tuning curves in the opposite direction (see example tuning curve). It is also known that each visual cortical layer has unique connections and these specific connections probably serve discrete functions. Thus, connections in each layer are dedicated for different purpose. Therefore, towards the same stimuli, neurons in different layers evoke different response properties indicating sequential stages of cortical processing (Martinez *et al.*, 2005). In summary, the application of sound differentially affected the layers 2/3 and 5/6.

### Layer 2/3 and layer 5/6 neurons change orientation preferences

In line with the previous studies in visual adaptation, many authors have described changes in orientation selectivity after a short (less than a minute) (Dragoi *et al.*, 2000) or long light adaptation (Ghisovan *et al.*, 2008; 2009; Bachatene *et al.*, 2012; Bachatene *et al.*, 2013; Bachatene *et al.*, 2015; Chanauria *et al.*, 2016) (several minutes). Usually, the protocol requires the frequent or continued application of a non-preferred property of the adapter such as orientation. In the above experiments, the peak of the orientation tuning curve is displaced either toward (attractive) or away (repulsive) from the adapting orientation. In all cases, such protocol results in the emergence of novel orientation selectivity. Neurons undergo a push-pull mechanism (Bachatene *et al.*, 2012; Bachatene *et al.*, 2013; Bachatene *et al.*, 2015; Chanauria *et al.*, 2016) that is characterized by the decrease and increase of the firing rates at the initially preferred and the adapting orientation (newly acquired) respectively.

In the current investigation, the non-preferred stimulus traditionally referred to as visual adapter is absent and is replaced by exposure to a sound stimulus imposed in seclusion for several minutes uninterruptedly. Results revealed that such presentation of sound shifts the orientation tuning curves of individual visual neurons in supra- and infra-granular layers of area 17 of cats. Consequently, the sound seemed to exert distinctive effects in addition to the global modulatory influence. As orientation selectivity is formed at the cortical level, thanks to aligned thalamic inputs upon the cortical recipient neurons, the changes of orientation selectivity are unlikely to happen at thalamic levels, where cells have mostly concentric receptive fields (although some units may exhibit a slight orientation bias). Further, we found populations of neurons (layer 2/3 and layer 5/6) exhibiting enhanced or diminished responses while the overall response of the population remained balanced (Figure 7A). Notably, at 90° orientation, a clear enhancement of response is seen whereas suppression occurs in layer 2/3 neurons. Layer 5/6 cells experienced an increase in excitation.

It has been noted that short duration sounds produce weak visual responses (Brady, 2011; Deneux *et al.*, 2018). Repeated presentation of the sound stimulus for a prolonged duration may be accountable for eliciting stronger responses in V1. Undoubtedly, the repeated stimulation aids in crossing a threshold capable of generating the required strength of interaction given that substantial auditory input reaches V1. This sort of mechanism can logically happen in layer 2/3 where pyramidal neurons receive direct inputs from the auditory cortex, as 12 min time is enough to encode the raw embedded visual information within the audio stimulus. The bias we observed may perhaps be a consequence of such a process, and that is why we notice an orientation bias towards horizontal axis for layer 2/3 neurons but not in layer 5/6. Also, this predisposition could be inherently induced because of the property of the sound itself. It has been known for many years now that visual cortex is a stimulus-driven structure (Doron *et al.*, 2002; Izraeli *et al.*, 2002; Piche *et al.*, 2007; Chabot *et al.*, 2008). The non-visual information in the stimulus can transmit from non-visual sensory structures to the visual cortex by direct cortico-occipital (temporo-occipital?) pathways and circumvent the higher order cortex. Xu and co-authors (Xu *et al.*, 2012) suggested that functional mixing of inputs from two different sources could allow for facilitative non-linear interactions within individual dendrites that may lead to the bias towards horizontal orientations. This non-linearity may yield a preference towards the horizontal orientations. Another important revelation from a previous study (Deneux *et al.*, 2018) suggests that the increasing intensity of sound strengthens the neuronal response. The intensity of the stimulus we used was kept the same for the 3s, and the same 3s time was repeated for 12 min.

### Inhibitory mechanisms

Recent studies have suggested that sound modulates light responses by impacting inhibitory neurons in the V1 cortex (Iurilli *et al.*, 2012; Kayser & Remedios, 2012; Pluta *et al.*, 2015; Ibrahim *et al.*, 2016). A very recent report (Deneux *et al.*, 2018) revealed that auditory cortex neurons project to V1 inhibitory neurons in superficial layers, especially layer 1. Indeed, area A1 sends axons to V1 where they connect to inhibitory interneurons (Iurilli *et al.*, 2012). As summarized by Meijer and authors in their paper (Meijer *et al.*, 2017), auditory stimuli suppress VIP interneurons which inhibit SOM cells which in turn suppresses PV (Parvalbumin; GABAergic interneurons) cells and distal dendrites (involved in non-local synapses) of supragranular pyramidal cells (Gentet, 2012; Meijer *et al.*, 2017). This specific pathway is activated precisely in response only to non-visual inputs (Deneux *et al.*, 2018). Indeed, pyramidal cells are entrenched by inhibitory cells (Tremblay *et al.*, 2016). Authors (Deneux *et al.*, 2018) reported that the observed inhibitory mechanisms originated in a dark setup, and these results could be applied to the current investigation since sound was applied in darkness. However, it must be underlined that inhibition was not directly tested in the present investigation consequently the above suggestions necessitate experimental evidence.

Furthermore, there is classical cross-orientation inhibition that may be added to above inhibitory processes initiated in A1 (Morrone *et al.*, 1982; Alitto & Dan, 2010; Priebe, 2016). Such steps may further contribute to changing of preferred orientation that result in shifts of selectivity. On similar lines, it has been hypothesized that sound exerts a divisive or additive influence, resulting in a decline or enhancement of light responses (Pluta *et al.*, 2015). These results were observed in mice yet similar explanations could apply to cats.

### Supramodal nature and cross-modal influences in the cortex

Our results demonstrate a new case of audio-visual communication, and further substantiate the observations made by previous investigations in cross-modal plasticity. Unlike the relatively slight modulation of firing rate of neurons detected in previous investigations in mice (Iurilli *et al.*, 2012; Ibrahim *et al.*, 2016), we observed a much more vigorous response-change in our data, wherein V1 neurons experienced a complete shift in their orientation preference. This kind of modification was reported earlier in literature related to adaptation studies where neurons were only exposed to an adapting orientation (visual stimulus) and acquired newly preferred selectivity. These visual adaptation protocols pertain to the classical flow of information involving LGN to the input layer 4 of the V1 and then to layer 2/3 and finally to layer 5/6. In the current case, the visual adapter was absent, yet similar results were obtained. Indeed, in this case, it is the application of sound for several minutes that induced neurons to acquire a new preferred orientation. Though the principle underlying such interactions has not been fully discovered yet, the most suitable explanation that fits in with our results is the combination of cross-modal interactions and the supramodal nature of the visual cortex. The visual cortex is a stimulus-driven structure. Therefore, it exhibits a unique response pattern towards specific protocols and stimuli. It is hypothesized that, upon sensory activation, the visual cortex relies on the abstract representation of the sensory stimuli regardless of the sensory modality. Stimulus-specific reorganization of the cortex can be responsible for the recalibration of the sensory system. In the present study, healthy animals were monocularly stimulated. A supportive study (Muckli *et al.*, 2015) highlights that a significant source of information lies in the context of the stimuli. Muckli and colleagues state that any sensory stimulus involving a feature can potentially embody the global features of the stimuli that are enough to trigger a complex scene representation of the same sound stimulus in the visual cortex (Bar, 2004; Oliva & Torralba, 2007), especially when presented for a prolonged duration. Although these effects have been discussed mainly from imaging studies on humans, the underlying principle supports the suggestion of our data (Vogels, 1999; Muckli *et al.*, 2005; Vetter *et al.*, 2014; Muckli *et al.*, 2015).

Besides, from the spectrogram it can be noticed that the sound stimulus has a structure and a direction in its frequency modulation. The stimulus quite creates the illusion of the oriented gratings for the auditory neurons and due to the repetitive nature of the stimulus it also has specific time-frequency organization that is yet again equivalent of oriented gratings for the auditory neurons. Therefore, these embedded features of the sound seem potential to modulate the activity of V1 neurons.

## Conclusion

The recent finding of the early involvement of the visual cortex in the integration of multimodal information has led to the testing of different stimuli towards non-corresponding primary sensory regions. During experimentation in this direction, different research groups have discovered a slight modulation of response with respect to short sound exposure. Our data demonstrate that prolonged sound presentation (sound adaptation) entirely modifies the tuning properties of individual cells and consequently reorganizes the cortical orientation maps in V1. Our results further substantiate and support that primary visual cortex is indeed inherently multisensory in nature. Nevertheless, a crucial and detailed inspection is required to fully fathom the mechanisms of communication between the primary sensory areas.

## Acknowledgments

NSERC Canada, Grant no 6943 supported this research. SM, NC, and VB designed the experiments. NC, VB, LB and SC participated in the experiments. NC wrote the paper and analyzed the data. JR greatly assisted with the optimization of the auditory stimulus, SM and JR contributed with comments that significantly improved the manuscript.

## References

Alitto, H.J. & Dan, Y. (2010) Function of inhibition in visual cortical processing. Curr Opin Neurobiol, 20, 340–346.

Atallah, B.V., Bruns, W., Carandini, M. & Scanziani, M. (2012) Parvalbumin-expressing interneurons linearly transform cortical responses to visual stimuli. Neuron, 73, 159–170.

Bachatene, L., Bharmauria, V., Cattan, S. & Molotchnikoff, S. (2013) Fluoxetine and serotonin facilitate attractive-adaptation-induced orientation plasticity in adult cat visual cortex. Eur J Neurosci, 38, 2065–2077.

Bachatene, L., Bharmauria, V., Cattan, S., Rouat, J. & Molotchnikoff, S. (2015) Reprogramming of orientation columns in visual cortex: a domino effect. Sci Rep, 5, 9436.

Bachatene, L., Bharmauria, V. & Molotchnikoff, S. (2012) Adaptation and Neuronal Network in Visual Cortex.

Bachatene, L., Bharmauria, V., Rouat, J. & Molotchnikoff, S. (2012) Adaptation-induced plasticity and spike waveforms in cat visual cortex. Neuroreport, 23, 88–92.

Bar, M. (2004) Visual objects in context. Nat Rev Neurosci, 5, 617–629.

Bardy, C., Huang, J.Y., Wang, C., FitzGibbon, T. & Dreher, B. (2006) ‘Simplification’ of responses of complex cells in cat striate cortex: suppressive surrounds and ‘feedback’ inactivation. The Journal of physiology, 574, 731–750.

Barlow, H.B., Blakemore, C. & Pettigrew, J.D. (1967) The neural mechanism of binocular depth discrimination. The Journal of physiology, 193, 327–342.

Barth, A.L. & Poulet, J.F. (2012) Experimental evidence for sparse firing in the neocortex. Trends Neurosci, 35, 345–355.

Bharmauria, V., Bachatene, L., Cattan, S., Brodeur, S., Chanauria, N., Rouat, J. & Molotchnikoff, S. (2016) Network-selectivity and stimulus-discrimination in the primary visual cortex: cell-assembly dynamics. Eur J Neurosci, 43, 204–219.

Bharmauria, V., Bachatene, L., Cattan, S., Chanauria, N., Rouat, J. & Molotchnikoff, S. (2015) Stimulus-dependent augmented gamma oscillatory activity between the functionally connected cortical neurons in the primary visual cortex. Eur J Neurosci, 41, 1587–1596.

Bouchard, M., Gillet, P.C., Shumikhina, S. & Molotchnikoff, S. (2008) Adaptation changes the spatial frequency tuning of adult cat visual cortex neurons. Exp Brain Res, 188, 289–303.

Brady, D. (2011) Mechanisms of Cross-Modal Refinement by Visual Experience.

Budinger, E., Heil, P., Hess, A. & Scheich, H. (2006) Multisensory processing via early cortical stages: Connections of the primary auditory cortical field with other sensory systems. Neuroscience, 143, 1065–1083.

Chabot, N., Charbonneau, V., Laramee, M.E., Tremblay, R., Boire, D. & Bronchti, G. (2008) Subcortical auditory input to the primary visual cortex in anophthalmic mice. Neuroscience letters, 433, 129–134.

Chanauria, N., Bharmauria, V., Bachatene, L., Cattan, S., Rouat, J. & Molotchnikoff, S. (2016) Comparative effects of adaptation on layers II-III and V-VI neurons in cat V1. Eur J Neurosci, 44, 3094–3104.

Deneux, T., Kempf, A. & Bathellier, B. (2018) Context-dependent signaling of coincident auditory and visual events in primary visual cortex. bioRxiv.

Denman, D.J. & Contreras, D. (2014) The structure of pairwise correlation in mouse primary visual cortex reveals functional organization in the absence of an orientation map. Cereb Cortex, 24, 2707–2720.

Doron, N.N., Ledoux, J.E. & Semple, M.N. (2002) Redefining the tonotopic core of rat auditory cortex: physiological evidence for a posterior field. The Journal of comparative neurology, 453, 345–360.

Dragoi, V., Rivadulla, C. & Sur, M. (2001) Foci of orientation plasticity in visual cortex. Nature, 411, 80–86.

Dragoi, V., Sharma, J. & Sur, M. (2000) Adaptation-induced plasticity of orientation tuning in adult visual cortex. Neuron, 28, 287–298.

Frenkel, M.Y., Sawtell, N.B., Diogo, A.C., Yoon, B., Neve, R.L. & Bear, M.F. (2006) Instructive effect of visual experience in mouse visual cortex. Neuron, 51, 339–349.

Fries, P., Nikolic, D. & Singer, W. (2007) The gamma cycle. Trends Neurosci, 30, 309–316.

Fritz, J., Shamma, S., Elhilali, M. & Klein, D. (2003) Rapid task-related plasticity of spectrotemporal receptive fields in primary auditory cortex. Nat Neurosci, 6, 1216–1223.

Gentet, L.J. (2012) Functional diversity of supragranular GABAergic neurons in the barrel cortex. Front Neural Circuits, 6, 52.

Ghazanfar, A.A. & Schroeder, C.E. (2006) Is neocortex essentially multisensory? Trends Cogn Sci, 10, 278–285.

Ghisovan, N., Nemri, A., Shumikhina, S. & Molotchnikoff, S. (2008) Visual cells remember earlier applied target: plasticity of orientation selectivity. PLoS One, 3, e3689.

Ghisovan, N., Nemri, A., Shumikhina, S. & Molotchnikoff, S. (2009) Long adaptation reveals mostly attractive shifts of orientation tuning in cat primary visual cortex. Neuroscience, 164, 1274–1283.

Hall, A.J. & Lomber, S.G. (2008) Auditory cortex projections target the peripheral field representation of primary visual cortex. Exp Brain Res, 190, 413–430.

Henry, G.H., Bishop, P.O., Tupper, R.M. & Dreher, B. (1973) Orientation specificity and response variability of cells in the striate cortex. Vision Res, 13, 1771–1779.

Ibrahim, L.A., Mesik, L., Ji, X.Y., Fang, Q., Li, H.F., Li, Y.T., Zingg, B., Zhang, L.I. & Tao, H.W. (2016) Cross-Modality Sharpening of Visual Cortical Processing through Layer-1-Mediated Inhibition and Disinhibition. Neuron, 89, 1031–1045.

Iurilli, G., Ghezzi, D., Olcese, U., Lassi, G., Nazzaro, C., Tonini, R., Tucci, V., Benfenati, F. & Medini, P. (2012) Sound-driven synaptic inhibition in primary visual cortex. Neuron, 73, 814–828.

Izraeli, R., Koay, G., Lamish, M., Heicklen-Klein, A.J., Heffner, H.E., Heffner, R.S. & Wollberg, Z. (2002) Cross-modal neuroplasticity in neonatally enucleated hamsters: structure, electrophysiology and behaviour. Eur J Neurosci, 15, 693–712.

Jeyabalaratnam, J., Bharmauria, V., Bachatene, L., Cattan, S., Angers, A. & Molotchnikoff, S. (2013) Adaptation shifts preferred orientation of tuning curve in the mouse visual cortex. PLoS One, 8, e64294.

Kayser, C. & Logothetis, N.K. (2007) Do early sensory cortices integrate cross-modal information? Brain Struct Funct, 212, 121–132.

Kayser, C. & Remedios, R. (2012) Suppressive competition: how sounds may cheat sight. Neuron, 73, 627–629.

Land, R., Engler, G., Kral, A. & Engel, A.K. (2012) Auditory evoked bursts in mouse visual cortex during isoflurane anesthesia. PLoS One, 7, e49855.

Liang, M., Mouraux, A., Hu, L. & Iannetti, G.D. (2013) Primary sensory cortices contain distinguishable spatial patterns of activity for each sense. Nat Commun, 4, 1979.

Liao, D.S., Krahe, T.E., Prusky, G.T., Medina, A.E. & Ramoa, A.S. (2004) Recovery of cortical binocularity and orientation selectivity after the critical period for ocular dominance plasticity. Journal of neurophysiology, 92, 2113–2121.

Lutcke, H., Margolis, D.J. & Helmchen, F. (2013) Steady or changing? Long-term monitoring of neuronal population activity. Trends Neurosci, 36, 375–384.

Maffei, L., Fiorentini, A. & Bisti, S. (1973) Neural correlate of perceptual adaptation to gratings. Science, 182, 1036–1038.

Martinez, L.M., Wang, Q., Reid, R.C., Pillai, C., Alonso, J.M., Sommer, F.T. & Hirsch, J.A. (2005) Receptive field structure varies with layer in the primary visual cortex. Nat Neurosci, 8, 372–379.

Meijer, G.T., Montijn, J.S., Pennartz, C.M.A. & Lansink, C.S. (2017) Audiovisual Modulation in Mouse Primary Visual Cortex Depends on Cross-Modal Stimulus Configuration and Congruency. J Neurosci, 37, 8783–8796.

Moore, B.D.t., Alitto, H.J. & Usrey, W.M. (2005) Orientation tuning, but not direction selectivity, is invariant to temporal frequency in primary visual cortex. Journal of neurophysiology, 94, 1336–1345.

Morrone, M.C., Burr, D.C. & Maffei, L. (1982) Functional implications of cross-orientation inhibition of cortical visual cells. I. Neurophysiological evidence. Proc R Soc Lond B Biol Sci, 216, 335–354.

Muckli, L., De Martino, F., Vizioli, L., Petro, L.S., Smith, F.W., Ugurbil, K., Goebel, R. & Yacoub, E. (2015) Contextual Feedback to Superficial Layers of V1. Curr Biol, 25, 2690–2695.

Muckli, L., Kohler, A., Kriegeskorte, N. & Singer, W. (2005) Primary visual cortex activity along the apparent-motion trace reflects illusory perception. PLoS Biol, 3, e265.

Muckli, L. & Petro, L.S. (2013) Network interactions: non-geniculate input to V1. Curr Opin Neurobiol, 23, 195–201.

Oliva, A. & Torralba, A. (2007) The role of context in object recognition. Trends Cogn Sci, 11, 520–527.

Piche, M., Chabot, N., Bronchti, G., Miceli, D., Lepore, F. & Guillemot, J.P. (2007) Auditory responses in the visual cortex of neonatally enucleated rats. Neuroscience, 145, 1144–1156.

Pluta, S., Naka, A., Veit, J., Telian, G., Yao, L., Hakim, R., Taylor, D. & Adesnik, H. (2015) A direct translaminar inhibitory circuit tunes cortical output. Nat Neurosci, 18, 1631–1640.

Priebe, N.J. (2016) Mechanisms of Orientation Selectivity in the Primary Visual Cortex. Annu Rev Vis Sci, 2, 85–107.

Ramoa, A.S., Mower, A.F., Liao, D. & Jafri, S.I. (2001) Suppression of cortical NMDA receptor function prevents development of orientation selectivity in the primary visual cortex. J Neurosci, 21, 4299–4309.

Ringach, D.L., Shapley, R.M. & Hawken, M.J. (2002) Orientation selectivity in macaque V1: diversity and laminar dependence. J Neurosci, 22, 5639–5651.

Schroeder, C.E. & Foxe, J.J. (2002) The timing and laminar profile of converging inputs to multisensory areas of the macaque neocortex. Brain Res Cogn Brain Res, 14, 187–198.

Stevenson, R.A., Wallace, M.T. & Altieri, N. (2014) The interaction between stimulus factors and cognitive factors during multisensory integration of audiovisual speech. Front Psychol, 5, 352.

Ten Oever, S., van Atteveldt, N. & Sack, A.T. (2015) Increased Stimulus Expectancy Triggers Low-frequency Phase Reset during Restricted Vigilance. J Cogn Neurosci, 27, 1811–1822.

Tremblay, R., Lee, S. & Rudy, B. (2016) GABAergic Interneurons in the Neocortex: From Cellular Properties to Circuits. Neuron, 91, 260–292.

van Atteveldt, N., Murray, M.M., Thut, G. & Schroeder, C.E. (2014) Multisensory integration: flexible use of general operations. Neuron, 81, 1240–1253.

Vetter, P., Smith, F.W. & Muckli, L. (2014) Decoding sound and imagery content in early visual cortex. Curr Biol, 24, 1256–1262.

Vogels, R. (1999) Categorization of complex visual images by rhesus monkeys. Part 2: single-cell study. Eur J Neurosci, 11, 1239–1255.

Xu, N.L., Harnett, M.T., Williams, S.R., Huber, D., O’Connor, D.H., Svoboda, K. & Magee, J.C. (2012) Nonlinear dendritic integration of sensory and motor input during an active sensing task. Nature, 492, 247–251.

